# Sex differences in DNA methylation in bats

**DOI:** 10.1101/2025.05.13.653776

**Authors:** Jack G. Rayner, Samantha L. Bock, Andrew J. Lonski, Nicole C. Riddle, Gerald S. Wilkinson

## Abstract

Sex-biased longevity is observed across a wide range of animal taxa, including bats, for reasons not well understood. Patterns of cytosine methylation vary predictably with age in many organisms, offering a valuable means to investigate differences in patterns of aging at the molecular level. We tested sex differences in cytosine methylation across 14 bat species and compared patterns of age-associated variation. Sex differences were overrepresented on the X chromosome, showing a strong pattern of female hypermethylation within promoter regions. Sex and age-associated differences in methylation were non-randomly distributed with respect to proximity to putative sex hormone receptor binding sites, with sites hypermethylated in males and females tending to be underrepresented near androgen and estrogen receptor binding sites, respectively. Across species, we observed the relative steepness of male versus female slopes of age-associated variation was associated with the strength of precopulatory sexual selection, with especially strong trends towards male-biased age-associated slopes in two harem-polygynous species that exhibit female-biased longevity. Our results offer insights into how patterns of methylation differ across sexes and ages, and raise intriguing questions for future research, such as whether sex differences in molecular aging reflect sex-biased longevity, for which records in bats are sparse.

## Introduction

The addition of methyl groups to cytosine nucleotides to form 5-methylcytosine (hereafter, ‘methylation’) is an important regulator of patterns of gene expression in many organisms ^1^, and rates of methylation at many CpG dinucleotides show predictable variation between tissues and across ages. Accordingly, epigenetic clocks have been developed that accurately estimate ages of individuals from various species ^2,3^, and for broader taxonomic groups ^4,5^, based on methylation profiles from a given tissue. Similar approaches have been used to predict biological sex from methylation data ^6,7^ which is particularly useful in species lacking sex chromosomes, such as those with environmental sex determination ^8,9^ and temperature-induced sex reversal ^10^. However, while sex and age represent factors of overarching interest in studies of genome-wide methylation patterns, their interaction (i.e., sex differences in methylation across ages) is less commonly considered. Moreover, sex chromosomes, which in addition to roles in biological sex determination appear to influence patterns of sex-specific aging ^11^, are often overlooked in studies of methylation ^12^.

Sex-biased longevity is observed across a broad range of taxa, but why and how these differences arise remains largely unclear ^13–15^. A possibility with some empirical support is that sex chromosome complement plays a direct role, with heterogametic individuals of many taxa appearing to suffer earlier mortality ^11^ perhaps due to deleterious consequences of sex-linked mutations ^16^, or the accumulation of transposable elements on degenerate, sex-limited chromosomes ^17^. Sex differences in life history, ubiquitous in sexually reproducing species, are also widely expected to play an important role. Differences between sexes in the cost of gamete production—sperm typically being much less costly to produce—are predicted to promote sex differences in resource allocation and life history ^18–20^. This pattern of anisogamy is thought to underlie a trend towards male-biased fitness benefits of multiple matings (i.e., Bateman gradients) in many species ^21^ and to generate strong male-male competition for mates, which is associated with selection for costly primary and secondary sexual traits ^22,23^. Investment in these traits may benefit reproductive fitness but involve allocating resources away from somatic maintenance, potentially accelerating male patterns of aging ^24^, consistent with the ‘disposable soma’ hypothesis ^25^.

In many organisms, sex differences in life history and resource allocation are regulated at least partly by the actions of hormones ^26^. Some have suggested that these hormones act to increase reproductive fitness by promoting growth during development but have deleterious consequences in later life, thereby accelerating patterns of aging ^27^—a form of antagonistic pleiotropy ^28^. In vertebrates, testosterone is consistently found to impair immune responses, while the effect of estrogen is more variable across different features of the immune response ^29^. Additionally, castration and the resulting reduction in androgen levels are associated with lifespan extension in mice and sheep ^30,31^, and a recent study found castration also slows epigenetic aging in sheep, based on differences between chronological age and age predicted by epigenetic clocks, potentially supporting cumulative effects of androgen exposure on male aging ^31^. By way of contrast, reduced expression of sex hormones in female mice and human women has been found to accelerate epigenetic aging, again determined by deviations between chronological age and that measures by epigenetic clocks ^32,33^, which could suggest female sex hormones reduce rates of aging. In line with these observations, estrogen is associated with anti-inflammatory and antioxidant activity ^34–36^. Altogether, existing evidence suggests sex hormones are likely to play important roles in influencing sex differences in molecular patterns of aging.

Bats are of particular interest in the study of aging ^37,38^, as they outlive most mammals of a similar size ^39^ despite high rates of infection by many disease-causing organisms ^40^. Many bat species are polygynous ^41,42^, promoting sex differences in life history which might influence patterns of aging and lifespan due to strong investment by males into reproduction-associated traits ^24^ and studies of sex steroid hormones in highly polygynous greater spear-nosed bats *Phyllostomus hastatus* and greater sac-winged bats *Saccopteryx bilineata* have shown androgens are elevated in harem males throughout much of the year ^43–45^. Recent studies of methylation in bats have revealed inter-specific differences in lifespan are reflected by patterns of methylation ^5^ and that epigenetic aging is slowed during hibernation^46^. However, sex differences in methylation of bat species have been largely uncharacterised.

In the current study, we compare patterns of methylation—a reliable aging biomarker ^47^— between sexes across 14 bat species, representing six families in the order Chiroptera. We identify sex-biased CpG sites and their representation across different genomic regions. We test for sex differences in slopes of age-associated methylation variation within each species, and test for an association between these differences and male testes size, which we use as a proxy measure inversely correlated with the opportunity for precopulatory sexual selection ^23,48–51^. Finally, we test for enrichment of sites showing sex-biased and age-associated methylation in regions nearby putative androgen and estrogen receptor binding sites. Our results provide insight into patterns of sex-biased methylation, identify patterns of sex-differences in the accumulation of methylation marks with age, and highlight the potential importance of sexually divergent life history strategies in shaping this variation.

## Material and Methods

### Data sources

Methylation data included in our analyses were obtained from Wilkinson et al. ^5^, who characterised patterns of age-associated change in methylation within and across bat species and developed epigenetic clocks to estimate individuals’ ages. Additional samples from *Eptesicus fuscus* were retrieved from Sullivan et al. ^46^, who characterised patterns of methylation associated with hibernation, though we excluded samples from hibernating bats. All data were collected from wing skin tissue samples which were assayed on the HorvathMammalMethylChip40 microarray, generating methylation values for 37,492 highly conserved CpG sites. We retrieved raw IDAT files for all samples from the NCBI GEO accession GSE164127. Then, we processed beta values (proportions of methylated vs. unmethylated sites) in SeSAMe v1.22.2 ^52^ using parameters recommended for the Mammal40 array platform (preprocessing code “SHCDPB”).

Prior to analysis, we excluded samples without known ages or with low confidence ages (=< 80 in ^5^). We excluded species lacking a minimum of five samples >= 0.3 years of age (at which point bats are likely to be fully grown) from males and females, each, retaining data from 14 species. We did not include samples from *Eidolon helvum* due to a lack of samples from older males. For *Molossus molossus*, we excluded data from a single very old female (5.8yrs), as there was no comparably old male from the same species. For *E. fuscus*, we excluded samples from one of two sources in the data from ^5^, due to strong skew in sex and age range that is not representative of the species in the wild. Finally, *P. hastatus* exhibit strongly sex-biased mortality, with males living up to about 11 years, while females can live up to 22 years ^43^. Because the maximum age of male *P. hastatus* in our sample was 5 years, approximately half of the maximum lifespan in wild populations, we excluded samples from females of this species older than 10 years of age, though doing so did not affect interpretation of any results. To our knowledge, maximum ages for samples of each sex of the remaining species are not obviously biased with respect to their relative life expectancies in wild populations (Fig. S1, Table S1).

### Methylation array annotations

Not all species present in our analyses have genome assemblies and/or annotations. For analyses requiring functional gene annotations and identification of X-linked sites we used an existing annotation for sites in the methylation array from *Rhinolophus ferrumequinum*, based on a chromosome-level genome assembly ^53^. We chose *R. ferrumequinum* as it is the bat species in which the greatest number (N=30,724) of CpG on the array can be mapped, with genes annotated ^5^. Most functional annotations are shared across bat species ^5^, and the X chromosome is highly conserved in bats as in other mammals ^54^, indicating this approach should not strongly bias our findings. Gene ontology-based analyses were performed by comparing genes of interest with the full list of genes in the array, using canine-based gene ontologies ^55^ (as canines are more closely related to bats than other well-annotated species such as humans or mice^53^), performed in the clusterProfiler R package ^56^ with a significance threshold of q < 0.05. Enrichr was used to identify associations between sex-biased or age-associated sites and putative transcription factor (TF) binding sites ^57^.

### Identification of sex-biased and age-associated sites

We identified sex and age differences in methylation using linear regression implemented using the DML function in SeSAMe v1.22.2. For each combination of site and species in our analysis, we ran the model *beta ∼ sex + age, w*here *beta* represents the proportion of methylated sequences for each sample at a given CpG site in the array and *age* is the z-scaled age (scaled across both sexes). We did not include a sex * age interaction term in our initial models, due to small sample sizes for some species. Instead, to detect sex differences in rates of age-related change at consistently age-associated sites, we ran sex-specific models of *beta ∼ age* in each species and compared distributions of sex-specific slopes.

### Testing sex differences in age-associated methylation change

To compare sex-specific rates of age-associated methylation, we identified in each species the subset of sites that showed evidence of age-associated changes (unadjusted-P < 0.05), in the same direction, in both sex-specific models (see above). We consider these to be sites which show at least some evidence of changing with age in the same direction in both sexes, between which it is reasonable to compare sex-specific slopes, whereas we did not consider sites that only showed evidence of age-associated differences in one sex, or which were age-associated in opposite directions. We removed sites for which predicted values in either sex, at any age, were within 0.05 of either boundary (0 or 1). For each site in this subset, we divided the male slope by the female slope and took the median value across sites for each species, and took the log_10_ of this ratio. We used both ordinary least squares (OLS) and phylogenetic generalized least squares (PGLS) regressions to test patterns of sex differences in slopes of age-associated methylation differences across species, with the latter accounting for phylogenetic non-independence of species in our sample. PGLS was implemented in the R package nlme (v 3.1-166) ^58^, against an assumption of Brownian evolution, though we confirmed that interpretation of our results did not differ when tested against other models (e.g., Pagel’s lambda implemented in caper v1.0.3 ^59^). Phylogenetic relationships were retrieved from TimeTree (accessed 25th July 2024) ^60^. We used OLS and PGLS to test for a significant deviation of the log_10_-transformed response variable, of median male divided by median female slopes across age-associated sites, from zero, which would indicate a consistent bias towards stronger slopes of age-associated methylation in one of the two sexes. Normality and heterogeneity of residuals was checked using diagnostic plots^61^.

An alternative method of comparing slopes of age-associated variation involves estimating relative rates of ‘epigenetic aging’, used in a variety of studies, by comparing known chronological ages with ages estimated from epigenetic clocks. In addition to the above approach, we also tested whether the accuracy of estimated ages differed notably between males and females of each species. We estimated ages of bats using an ‘all-bat’ epigenetic clock, which is designed to estimate chronological age for any species of bat based on methylation profiles in skin samples ^5^. We then ran a linear regression of the clock-estimated age against known chronological age for each species and compared residuals between males and females.

### Testing associations between age-associated methylation and male reproductive investment

Next, we tested whether sex-differences in the slope of age-associated methylation variation showed an association with inferred differences in the strength of precopulatory sexual selection. Specifically, we used relative male testes mass for each species as a proxy measure of sperm competition, the risk of which is strongly reduced when males can monopolise access to females (e.g., in harem polygynous mating systems) ^51^. Consequently, males of species with strongly polygynous-skewed mating systems are expected to invest more heavily in traits under precopulatory sexual selection, such as those involved in male-male competition, rather than sperm competition. This apparent tradeoff between investment in pre- and postcopulatory sexual selection is observed in bats ^23,48^, pinnipeds ^50^, anurans ^62^ and cetaceans ^49^.

We obtained measures of relative testes and body mass for several species from ^48^ and ^41^. For *Pteropus hypomelanus,* and *P. vampyrus*, we retrieved measures from ^63^. Data for *Eptesicus fuscus* was retrieved from the Division of Mammals Collections at the Smithsonian Museum of Natural History ^64^. Data for *P. pumilus* were provided by DeeAnn Reeder, who collected the data for a previous study ^65^. In each case, testes mass was extrapolated from testis length and width measurements (mm), used to calculate the volume of a prolate spheroid (0.5236 x length x width^2^) ^48^. To calculate relative testes mass, this volume was doubled and assumed to have the same weight as water, then divided by average male body mass. Data from *Molossus molossus* were retrieved from ^66^, in which body weight and testes weight were measured directly. We used PGLS to test for a statistical association between male-biased slopes of age-associated methylation variation and the relative testes mass in each species, each log_10_-transformed. We predicted that species in which males have smaller relative testes mass, which we presume to be under stronger precopulatory sexual selection, would show a stronger bias towards male-biased slopes.

A caveat to the above analyses is that slopes of age-associated variation might be influenced by differences in sample size and, in particular, age distribution between sexes, which were evident for some of the species in our sample—in some cases due to sex-biased longevity (Table S1, Fig. S1). To address this to the best of our ability, we investigated whether the interpretation of results was affected if we: 1) equalized age ranges by removing samples from individuals aged 1 or more years younger/older than the youngest/oldest individual of the opposite sex; or, 2) equalized age distributions and sample sizes by retaining only pairs of males and females of each species that were aged within 1.5yrs of one another. The latter analysis required we drop the *Tadarida brasiliensis* species from our analysis as only 2 samples of each sex remained.

### Tests for involvement of sex hormones

Androgen sensitive CpG sites on the HorvathMammalMethylChip40 microarray were defined previously by ^31^ as sites exhibiting differential CpG methylation in response to castration and the consequent reduced androgen production in domesticated sheep, a subset of which were shown to also overlap with known human targets of androgen receptor (AR) based on ChIP-seq data. As a first test for a conserved role of androgens in regulating sex-biased methylation patterns in bats, we tested overlap between these previously identified androgen-sensitive sites and sites exhibiting sex-biased methylation in the present study.

As a complementary approach to understand the potential role of sex steroid hormones in regulating sex-biased and sex-specific age-associated methylation patterns, we performed a genome-wide scan of the *R. ferrumequinum* genome for putative binding sites of AR, estrogen receptor alpha (ESR1), and estrogen receptor beta (ESR2) using the FIMO (Find Individual Motif Occurrence) tool within the MEME suite (v 5.5.5) ^67^. We derived hormone receptor binding motifs from the JASPAR 2024 database (MA0007.2 for AR, MA0112.1 for ESR1, MA0258.1 for ESR2) ^68^ in MEME Motif Format. Putative binding sites were defined according to a significance threshold of P < 3e-7, yielding 17,502 AR binding sites, 13,110 ESR1 binding sites, and 5,942 ESR2 binding sites. We quantified the proximity of each CpG site in *R. ferrumequinum* to the nearest putative binding site using the GenomicRanges package (v. 1.56.1) in R ^69^ and defined sites according to their location within 5 kb of the nearest binding site. To extend annotations of putative binding sites identified in *R. ferrumequinum* to the other species included in our sex-biased analysis, we filtered sites to only include those which mapped to and had the same nearest gene in the *R. ferrumequinum* (member of Yinpterochiroptera suborder) genome and in the genome of at least one species of *E. fuscus*, *D. rotundus*, *P. discolor*, *P. hastatus*, or *S. bilineata*, from the Yangochiroptera suborder. We tested whether unique sex-biased sites (P_adj._ < 0.05) identified across species were over- or underrepresented in genomic regions proximate to putative hormone receptor binding sites with Fisher’s exact tests. For the purposes of this analysis, sites showing male-biased and female-biased methylation were analysed separately and defined according to the directionality of methylation difference in the majority of species. Sites for which there were equal numbers of species exhibiting male- and female-biased methylation were excluded from the analysis. We also examined the representation of sites exhibiting sex-specific age-associated methylation patterns in regions proximate to binding sites. All unique sites that met the criteria above and showed evidence of age-associated methylation (unadjusted P < 0.05) in both sexes in concordant directions were included.

R v.4.4.1 ^70^ was used to perform data analysis. Additional results files, metadata, and scripts underlying analyses are available at https://osf.io/v6c5n/. Raw methylation data were previously made available from the NCBI GEO under accession GSE164127^5,46^.

## Results

Below, we present results from analyses of methylation data across 30,724 CpG sites from 403 bat skin samples (249 female, 154 male) across 14 species, with broad representation across the Chiroptera (eight species from the suborder Yangochiroptera, six from the Yinpterochiroptera). (Table S1; Fig. S1)

### Patterns of sex-biased methylation

Patterns of sex-biased methylation from the full model (*beta ∼ sex + age*) largely fit expectations by strongly highlighting the X chromosome. Across species, 2,108 unique sites (6.86% of the sites mapped to the *Rhinolophus ferrumequinum* genome) showed significant (P_adj_ < 0.05) sex-biased methylation in one or more species, compared with 5,816 sites (18.93%) showing significant age-associated variation in any species. These sex-biased sites were massively overrepresented on chromosome 1 (syntenic with the human X chromosome) (Fig 1A,B) (Fisher’s exact test: odds ratio = 227.960, P < 2.2e-16). We observed hypermethylation in both sexes across sex-biased sites, though there was an overwhelming pattern of female hypermethylation in X-linked promoter regions (Fig. 1B). As methylation of promoter regions is thought to often suppress gene expression ^1^, this pattern might be associated with dosage compensation through female X-inactivation. Sites showing sex-biased methylation were overrepresented in exon regions of six species (Fisher’s exact test: P_adj_ < 0.05), with no other genome features significantly overrepresented in any species (Fig 1C).

**Figure 1.**
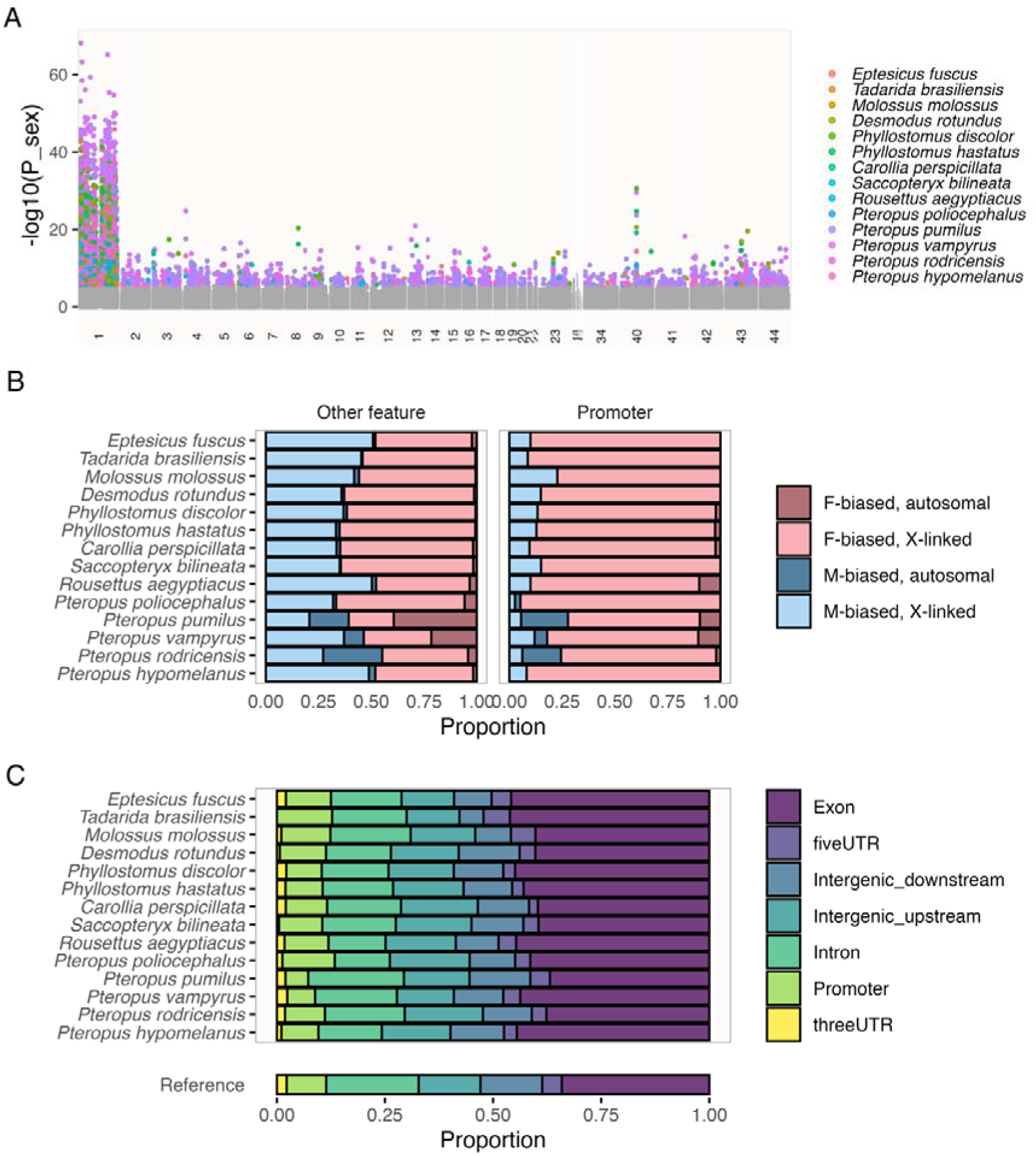
Sex-biased patterns of CpG methylation. **A**) Genome-wide distribution of significant (coloured by species) sex-biased sites, highlighting involvement of Chromosome 1 (syntenic with the human X chromosome). Chromosome identifiers are shown on the X-axis, and the log10 of the P-value for each site is shown on the Y-axis. **B**) Proportions of sites showing significant sex-biased methylation in each species, coloured by direction of sex-bias (i.e. which sex shows the higher level of methylation) and X/autosomal linkage, and whether present in promoter regions. **C**) Proportions of sites showing significantly sex-biased methylation levels within different types of genomic features.

In contrast with sites showing sex-biased methylation, age-associated sites identified in the full model were significantly underrepresented on the X chromosome; X-linked sites were 0.813 times as likely as autosomal sites to show age-associated differences in methylation (Fisher’s exact test: P = 0.014) consistent with a previous analysis ^5^. Among age-associated sites, promoter regions were significantly overrepresented in six species—replicating results from ^5^, and as observed in dogs by ^71^—while upstream intergenic and 5’ UTR regions were each overrepresented in two species. For a full discussion of age-associated patterns of methylation in bats, see: Wilkinson et al. ^5^.

Next, we investigated patterns of sex-biased and age-associated methylation across species, and compared these between X-linked and autosomal sites. On autosomes, very few CpG sites with sex-biased methylation in one species showed consistent, significant male- or female-biased methylation in other species (Fig. 2A). Notably, a large proportion of these autosomal sites were significantly sex-biased in species from the family *Pteropodidae* (Fig. 1B). In contrast, there was much greater conservation of sex-biased CpG methylation patterns on the X chromosome, perhaps owing to shared patterns of dosage compensation through X inactivation (Fig. 2A), and 29 cytosines showed sex-biased methylation in all 14 species. Of 280 sites with sex-biased methylation in the majority (> 7) of species tested, 190 were consistently hypermethylated in females, whereas 90 were consistently hypermethylated in males. Lists of genes associated with consistent male or female-biased methylation showed no overrepresentation of any gene ontology terms at P_adj_ < 0.05, though sites showing sex-bias in at least one species reported overrepresentation of biological processes involved in metabolic and RNA synthesis, and nervous system development (Fig. S2). Interestingly, lists of annotated genes associated with CpG sites showing consistently male-biased or female-biased methylation were enriched for genes regulated by the same five transcription factors: PRDM5, perhaps notable for its apparent role in tumour suppression^72^; E2F1, which plays an important role in apoptosis ^73^; TFAP2A; RUNX2, associated with skeletal development and adult bone density ^74^, and SOX2. Genes nearby CpG sites hypermethylated in females were also significantly associated with POU5F1 and GLI1. SOX2^75^ and POU5F1 ^76^ are pluripotency factors and were previously found to be associated with clusters of CpG sites that show consistent changes with age across mammals ^77^. Finally, a site located on Chr40 (Fig 1A) that showed female-biased methylation in 13 species was annotated for proximity to POU3F2, a transcription factor involved in neuronal differentiation ^78^.

**Figure 2.**
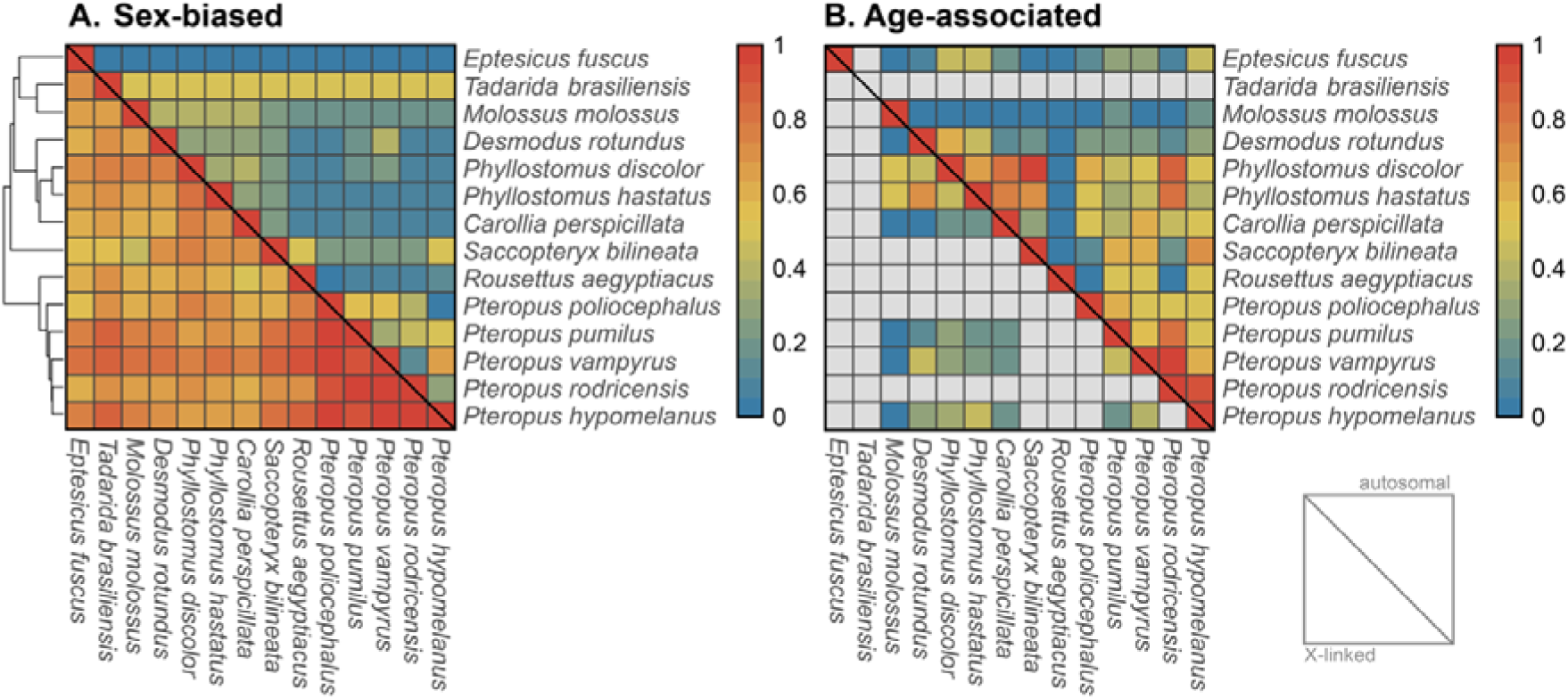
Cross-species overlap in sex-biased and age-associated CpG sites. Proportions of cytosines showing concordant directions of A) sex-biased and B) age-associated variation in methylation levels shared across species. The upper-right diagonal shows comparisons between autosomal CpG sites, and the lower-left diagonal comparisons between X-linked sites. Grey cells are those with no data (i.e., no age-associated sites in one of two species being compared). Rows and columns are ordered according to the species cladogram (far left).

### Sex differences in age-associated methylation

We next investigated whether and how patterns of age-associated variation appeared to differ between males and females across species. First, we tested for sex differences in age-associated methylation patterns with respect to their genomic locations and putative functions of nearby genes. We ran sex-specific analyses of age-associated variation (*beta ∼ age*), for each species. For each species, sex-specific slopes of age-associated change across sites were strongly correlated across CpG sites (Spearman’s rank correlation: mean rho = 0.406; all P < 1e-161) (Fig. S4). Interestingly, however, these correlations were significantly stronger among autosomal than X-linked sites (Wilcoxon signed rank test: P < 0.001). We also found that CpG sites that were significantly age-associated in either sex of any species at Padj < 0.05 were not significantly more or less likely to be located on the X-chromosome than predicted by chance (Fisher’s exact test: P > 0.10). This contrasts with the finding above that age-associated sites identified by the full model were underrepresented on the X, and could indicate that sites on the X chromosome are subject to sex x age interactions ^79^.

Next, we investigated CpG sites showing evidence of age-associated differences in methylation, in the same direction, in both sexes (P_unadj._ < 0.05 in both). We divided each male slope by the corresponding female slope, and took the log_10_-transformed median of this ratio for each species as a measure of the relative male slope of age-associated variation.

Ordinary least squares regression reported that this log_10_-transformed ratio differed from 0 (F_1,13_ = 9.283, P = 0.009) with males having steeper slopes indicative of stronger age-associated variation, albeit with exceptions in three species of the *Pteropodidae*: *Pteropus vampyrus*, *Pteropus poliocephalus*, and *Rousettus aegyptiacus* (Fig. 3). Consistent patterns of male-biased slopes were observed among X-linked and autosomal sites (Fig. S5-S6), and if we adjusted for sex differences in age range and sample sizes (both P < 0.05; Fig. S7-8) (see Methods). However, phylogenetic generalized least squares regression (PGLS) revealed patterns of male-biased slopes of age-associated variation were not significant once phylogenetic non-independence of species was accounted for (F_1,13_ = 1.342, P = 0.268). We also did not find any indication that our observation of male-biased slopes of change at age-associated sites was reflected by deviations between estimated and known chronological ages, observing no evidence of any sex differences in residuals from regression of estimated age on known age across species (F_1,13_ = 0.292, P = 0.598) (Fig. S9).

**Figure 3.**
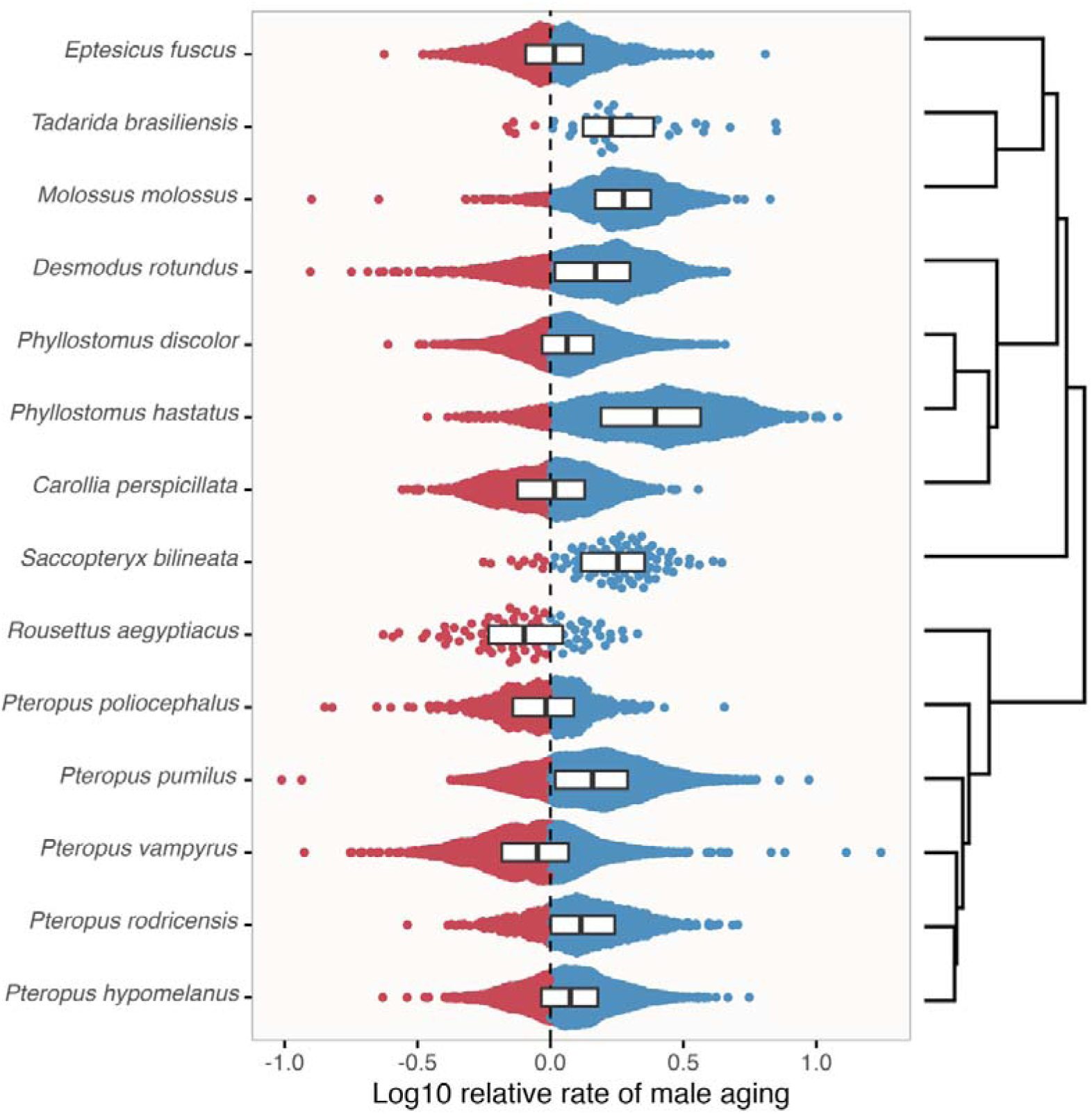
Sex-biased rates of change at age-associated CpG sites. Each point illustrates relative rates of change in the proportion of methylated CpG sequences with age among sites that showed evidence of correlated age-associated change (P_unadj._ < 0.05) in both sexes in each species. Note the log10 scale, such that values greater-than or less-than 0 (blue and red, respectively) represent male and female-biased slopes of age-associated variation, respectively. Species are shown on the Y-axis, with the corresponding phylogeny shown to the right of the plot.

We next tested for an association, across species, between measures of the relative male slope of age-associated variation (i.e., the medians illustrated by boxplots in Fig. 3), and relative testes mass: a proxy of the risk of sperm competition, which we treat as indicative of weaker precopulatory sexual selection. Log_10_ relative testes mass was significantly negatively associated with relative male slope after accounting for phylogenetic non-independence of species (PGLS: F_1,12_ = 8.963, P = 0.011) (Fig. 4). This general pattern remained after subsampling procedures designed to adjust for sex differences in age distribution and/or sample size, specifically: 1) removal of samples from individuals aged 1 or more years younger/older than the youngest/oldest individual of the opposite sex (Fig. S10) and, 2) retaining only pairs of males and females of each species that were aged within 1.5yrs of one another (Fig. S11). However, in the former analysis the pattern was not significant at P < 0.05 (P = 0.065), while results of the latter analysis did not confirm to PGLS assumptions of normally distributed residuals.

**Figure 4.**
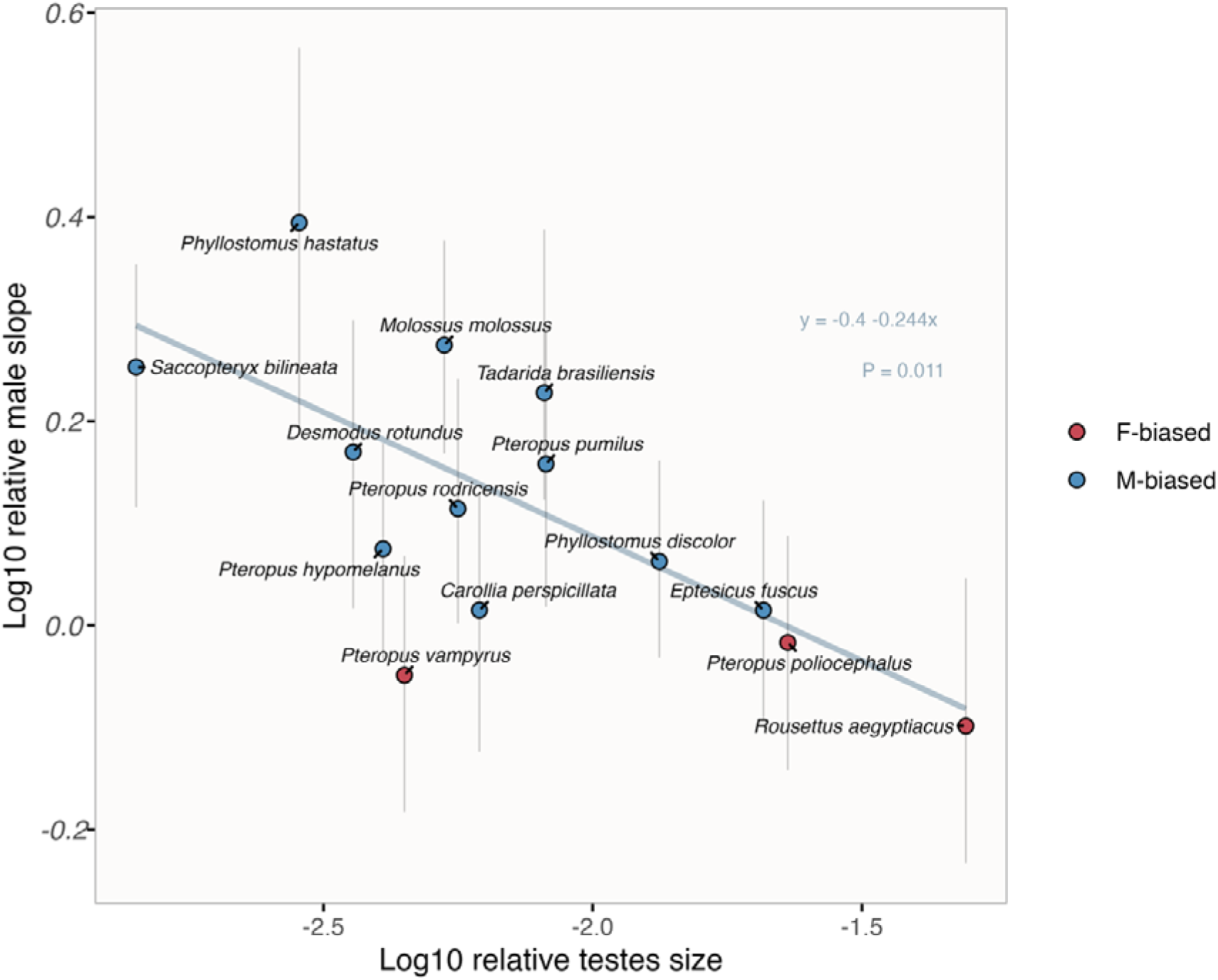
Relationship between relative testes mass and relative rates of methylation change in males. Points and vertical lines illustrate median values and interquartile ranges with respect to relative male slopes of age-associated variation, also shown in Fig. 3. Blue and red points represent species in which males and females tended to have stronger slopes of age-associated variation, respectively. The solid line visualises the relationship predicted by phylogenetic generalized least squares regression.

### Involvement of sex steroid hormones

Finally, we tested the putative involvement of sex steroid hormones in contributing to sex and age differences in methylation we observe. Consistent with our expectation, androgen-sensitive CpG sites previously identified by ^31^ in sheep were more likely to be reported as sex-biased in the present study, but the effect was relatively minor (odds ratio = 1.26; Fisher’s exact test: P < 0.001). Of the 4,092 sites in our analysis that were identified as androgen-sensitive by ^31^, 339 (7.92%) reported significantly sex-biased methylation in any bat species. Similarly, among age-associated CpG sites, there was significant overrepresentation of androgen-sensitive sites (odds ratio: 1.328, P < 0.001). Sugrue et al. ^31^ noted with particular interest that an androgen-sensitive site on the HorvathMammalMethylChip40 array, cg21524116 located within the MKLN1 gene in sheep, showed consistent elevated rates of age-associated change in CpG methylation in males of sheep, mice, and bats. However, this CpG site is not annotated for proximity to any genes in the *R. ferrumequinum* genome, so it was not included in our analyses. Nevertheless, two other sites (cg09596855 and cg17030869) were annotated for proximity to MKLN1 in *R. ferrumequinum*, and cg17030869 showed evidence of male-biased patterns of methylation change with age (P < 0.05 in males, with male-biased slope of change) in *P. hastatus* and *S. bilineata*, two species with female-biased longevity and strongly male-biased age-associated slopes (Fig. 2).

We next investigated under/overrepresentation of CpG sites with male-biased or female-biased methylation in regions proximate to putative androgen receptor (AR) and estrogen receptor binding sites identified in the *R. ferrumequinum* genome. Sites with male-biased methylation were significantly enriched in regions proximate (< 5 kb) to putative estrogen receptor alpha (ESR1) (odds ratio = 1.82; Fisher’s exact test: P < 0.001) and estrogen receptor beta (ESR2) binding sites (odds ratio = 2.36; P < 0.001) but underrepresented near AR binding sites (odds ratio = 0.66; P = 0.047). By contrast, CpG sites with female-biased hypermethylation were non-significantly underrepresented in regions proximate to putative ESR1 (odds ratio = 0.79; P = 0.069) and significantly underrepresented near ESR2 binding sites (odds ratio = 0.58; P = 0.016), but did not differ from expectation near AR binding sites. (Fig. 4A)

CpG sites exhibiting a higher rate of age-associated change in males were significantly underrepresented in regions proximate to putative AR binding sites identified in the *R. ferrumequinum* genome (odds ratio = 0.8, P < 0.001), significantly overrepresented near ESR1 (odds ratio = 1.21, P < 0.001), and non-significantly overrepresented near ESR2 binding sites (odds ratio = 1.15, P = 0.080). Surprisingly, CpG sites exhibiting a greater slope of age-associated variation in females were also underrepresented near AR binding sites (odds ratio = 0.84, P = 0.039), but did not differ from expectation near ESR binding sites. (Fig. 4B)

**Figure 4.**
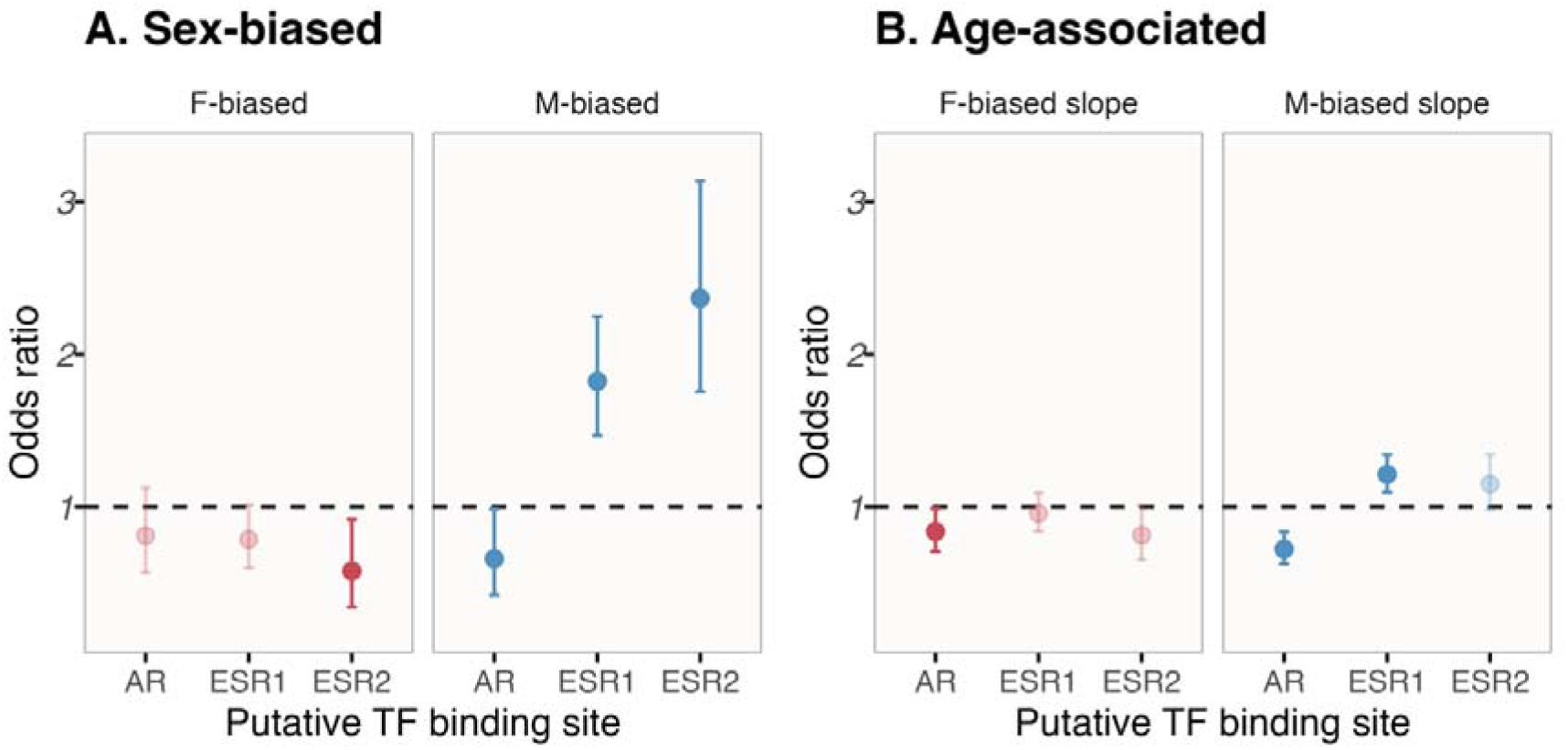
Results of tests for over or underrepresentation of sites showing sex-biased methylation and sex-biased slopes of methylation change with age within 5 kb of putative binding sites for androgen and estrogen-associated transcription factors. Error bars show 95% confidence intervals, with darker points those reported as significantly over- or underrepresented at P_unadj._ < 0.05.

## Discussion

Factors that might underpin patterns of sex-biased aging and/or mortality have attracted substantial recent interest^11,13,14^, but their respective contributions remain unclear. Our results provide insights into sex differences in methylation in general and across ages, revealing that a substantial portion of CpG sites show sex-biased patterns of methylation. We also identified a general pattern of stronger age-related variation in methylation among males of each species, albeit one that was not significant after accounting for phylogenetic non-independence, but that appeared to be associated with species differences in the inferred strength of precopulatory sexual selection. An open question in the study of sex-biased longevity is whether differences in lifespan are reflected by different rates of molecular or physiological aging^15^, and we observed patterns of male-biased methylation change in two species, *Phyllostomus hastatus* and *Saccopteryx bilineata*, in which female-biased longevity has been repeatedly observed ^43,80,81^.

Perhaps unsurprisingly, we found sex differences in methylation were strongly overrepresented on the X chromosome. When considering X-linked sites located in promoter regions, within which methylation is widely expected to suppress gene expression ^1^, we observed a strong pattern of female-biased methylation that was consistent across species. In mammals, sex chromosome dosage compensation is observed to occur via female inactivation of a single copy of the X, though studies in mice and humans have shown variable proportions of the X chromosome escape X-inactivation ^82^. Thus, the pattern of female-biased methylation in X-linked promoter regions seems likely to be associated with conserved patterns of X-inactivation across bat species. In contrast, we did not observe enrichment of the X chromosome among age-associated sites across models including both sexes, or sex-specific models. However, studies of humans and dogs have indicated sites that differ in age-associated patterns of methylation between sexes are enriched on the X chromosome ^71,83^, and we found evidence supporting this in the weaker correlation of sex-specific slopes among age-associated CpG sites on the X chromosome.

Across CpG sites that showed evidence of age-associated variation in both sexes, we observed that males exhibited stronger patterns of age-related differences in 11 of the 14 bat species in our study, though an important caveat is that this trend was not significant after accounting for phylogenetic non-independence of the species in our analysis. While X chromosomes did not show pronounced age-related changes in methylation, differences in sex chromosome complement might nevertheless influence sex-differences in rates of physiological aging, and are widely expected to do so^14^. For example, the ‘unguarded X’ hypothesis suggests that heterogametic individuals will suffer stronger fitness consequences of harmful X-linked alleles ^84^. The ‘toxic Y’ hypothesis, on the other hand, predicts that harmful mutations and, in particular, repetitive elements will accumulate on non-recombining sex chromosomes (i.e., Y or W chromosomes) with deleterious consequences ^85^. In either case, heterogametic individuals will be disadvantaged, potentially accelerating patterns of aging and/or reducing their lifespan. Consistent with this, a trend towards reduced lifespan in the heterogametic sex is observed across several animal taxa, including tetrapods and insects ^11,86^, and across birds and mammals in which females typically exhibit similar sex roles but are hetero- and homogametic, respectively ^87^. Since the bats in our study follow the typical mammalian XX/XY sex-determination system, or XX/XY_1_Y_2_ in *Carollia perspicillata* ^88^, the steeper slopes of age-associated variation in methylation we observe in males could be influenced by toxic Y and/or unguarded X effects. However, recent work has disputed whether unguarded X or toxic Y effects are sufficiently strong to explain sex differences in longevity ^16,89^.

We observed the trend towards male-biased slopes of age-associated variation was variable across species, and had notable exceptions. This cross-species variation could support alternative explanations of sex-differences in aging such as the disposable soma theory of aging ^25^, in which strong investment in reproduction-associated traits is predicted to trade-off against somatic maintenance. We found some support for this idea, in that the relative magnitude of male-biased age-associated methylation variation was positively associated with the inferred strength of precopulatory sexual selection across species, based on the presumption of an inverse correlation between precopulatory sexual selection and relative testes size. Several of the bats in our sample exhibit harem polygynous mating systems ^42,48^, in which males are predicted to invest heavily in secondary sexual traits such as body size ^90^, and a broader taxonomic sample may convincingly demonstrate whether sex-biased rates of age-associated methylation variation differ predictably based on mating system. Accelerated ‘epigenetic aging’ associated with reproductive investment has been observed in sheep and humans ^3,91^, but we are unaware of a cross-species analysis.

Sex hormones play an important role in regulating resource allocation towards reproduction versus somatic maintenance ^26^. In both *Phyllostomus hastatus* and *Saccopteryx bilineata*— two harem polygynous bat species with female-biased longevity and greater slopes of age-associated methylation change in males—harem males have high levels of testosterone throughout much of the year ^43,45,80^. Given this observation, and the hypothesised link between androgens and male aging, we might expect sites showing male-biased slopes of age-associated variation in methylation to be enriched in regions of the genome associated with regulation by androgens. Previously identified androgen-sensitive sites in sheep ^31^ were overrepresented among sex-biased and age-associated sites in the present study. Moreover, in genomic regions proximate to putative hormone receptor binding sites, we observed that CpG sites showing male-biased and female-biased methylation patterns were over- and underrepresented, respectively, at putative estrogen receptor binding sites. This pattern is broadly consistent with expectations, based on sex differences in endocrine regulation, if it is assumed that methylation is predominantly associated with suppression of gene expression. Next, sites exhibiting more pronounced age-associated variation in males were strongly underrepresented in regions proximate to putative androgen receptor binding sites, but were generally overrepresented near putative estrogen receptor binding sites. This raises the possibility that genomic regions directly regulated by the DNA binding activity of the androgen receptor may be less susceptible to age-associated methylation change in males. This pattern could feasibly relate to changes in social dominance with age; for example, older male *P. hastatus* are more likely to attain harem status ^43^, would could select against age-associated changes in methylation at androgen-associated loci. In general, results presented here and in other mammals ^3,31,92^ underscore the importance of future work examining the mechanisms by which androgens and other sex steroid hormones interact with the aging epigenome in species representing diverse mating systems.

Methods of estimating individuals’ ages based on methylation profiles are becoming increasingly popular, with epigenetic clocks having been developed for a wide variety of organisms. Though their application is relatively costly in most organisms (but see ^93^), these techniques provide powerful tools in studies of wild animals, in particular, when age records might otherwise involve labour-intensive observation and/or destructive sampling. Another common application of these clocks, particularly in humans, is to quantify accelerated epigenetic or ‘biological’ aging based on deviations between ages estimated from methylation profiles and known chronological ages, with individuals whose age is overestimated considered to exhibit accelerated epigenetic age ^3,31,46,94^. In our analysis, we did not observe an association between these deviations and sex across species. This clock-based approach appears to sometimes work well when ‘aging clocks’ have been specifically developed to incorporate morbidity and other factors to estimate lifespan or physiological aging ^94–96^. However, the broader applicability of this approach, particularly the application of clocks that are designed to estimate chronological age, is perhaps less clear^97^. For example, Kabacik et al. ^98^ found little evidence of an association between epigenetic age and biomarkers of senescence in primary human cell cultures, suggesting age estimates better reflected chronological rather than biological age. In fact, it has been noted that improved accuracy in predicting chronological age necessarily trades off with accuracy in predicting mortality and associated factors ^94,99^. Overall, we believe the fact that our findings of stronger age-related differences in methylation among males were not recapitulated by the ‘epigenetic age’ analysis most likely reflects this clock having been built specifically to accurately estimate chronological age of bats from skin samples.

## Conclusion

Our result shed light on sex differences in methylation and age-related changes across several bat species. Sex-biased patterns of methylation were common across species and were strongly enriched on the X-chromosome. Sex-biased methylation patterns of many X-linked CpG were shared across multiple species, which may be associated with conserved mechanisms of dosage compensation in females. We observed stronger slopes of age-related variation in methylation among males in several species, though the pattern was not considered statistically significant after accounting for their phylogenetic non-independence. Whether this trend might be associated with sex-differences in lifespan is an intriguing question with some equivocal support from our limited sample, and we also found some support for an association between male-biased slopes of age-associated variation and the inferred strength of precopulatory sexual selection acting on males. Sex and age-associated differences in methylation were non-randomly distributed with respect to sex hormone receptor binding sites, with CpG sites showing hypermethylation in males and females tending to be underrepresented in regions nearby putative androgen and estrogen receptor binding sites, respectively. Two limitations of our study are that we rely on cross-sectional samples of DNA methylation to infer age-associated changes, and that some species had sex-skewed age or sample distributions that could influence age comparisons. Future research incorporating longitudinal sampling of similar numbers of males and females of multiple species is likely to offer increased resolution in understanding how sexes differ in trajectories of molecular aging, and whether this contributes to sex differences in longevity.

## Acknowledgements

We thank DeeAnn Reeder for sharing with us testes measurement data from *P. pumilus*. This work was supported by the National Science Foundation under Award No. DBI-2213824 to NCR and GSW, and the National Institutes of Health under Award No. R61-AG078474 to GSW.

## Author contribution

**Conceptualization**: JGR, GSW. **Formal analysis**: JGR, SLB. **Funding acquisition**: NCR, GSW. **Writing – original draft**: JGR, SLB. **Writing – review and editing**: all authors.

## Supporting Information

**Table S1.** Numbers and age distributions of samples from each species included in the analysis. Numbers in parentheses represent values before removal of samples < 0.3 years of age. Numbers of sex-biased sites, and sites showing evidence of age-associated differences in both sexes, are also given.

| Species | Sex | N | Min. age | Max. age | Sex-biased sites<sup>a</sup> | Age-associated<sup>b</sup> |
| --- | --- | --- | --- | --- | --- | --- |
| *Carollia perspicillata* | F | 16 | 1.1 | 9.8 | 439 | 1719 |
|  | M | 13 | 0.5 | 10.5 |  |  |
| *Desmodus rotundus* | F | 25 | 0.3 | 16.1 | 440 | 1574 |
|  | M | 15 | 1 | 17.3 |  |  |
| *Eptesicus fuscus* | F | 9 | 1.72 | 7.1 | 288 | 1027 |
|  | M | 9 | 0.72 | 7.1 |  |  |
| *Molossus molossus* | F | 8 | 0.3 | 3.7 | 194 | 1605 |
|  | M | 5 | 0.3 | 1.7 |  |  |
| *Phyllostomus discolor* | F | 25 | 0.3 | 17.7 | 550 | 5326 |
|  | M | 14 | 0.3 | 12.2 |  |  |
| *Phyllostomus hastatus* | F | 33 | 1 | 10 | 454 | 1393 |
|  | M | 5 | 1.8 | 5 |  |  |
| *Pteropus hypomelanus* | F | 29 | 0.4 | 19.3 | 573 | 2825 |
|  | M | 12 | 0.4 | 19 |  |  |
| *Pteropus poliocephalus* | F | 10 | 7.2 | 16.7 | 314 | 441 |
|  | M | 6 | 6.1 | 15.8 |  |  |
| *Pteropus pumilus* | F | 24 | 1.6 | 16.9 | 1365 | 3169 |
|  | M | 22 | 0.8 | 17.3 |  |  |
| *Pteropus rodricensis* | F | 12 | 7 | 19.3 | 531 | 701 |
|  | M | 7 | 4 | 20.9 |  |  |
| *Pteropus vampyrus* | F | 27 | 0.6 | 22.4 | 1041 | 3270 |
|  | M | 24 | 1 | 20.7 |  |  |
| *Rousettus aegyptiacus* | F | 8 | 5 | 10.1 | 394 | 85 |
|  | M | 8 | 7 | 14 |  |  |
| *Saccopteryx bilineata* | F | 18 | 0.3 | 8.3 | 200 | 86 |
|  | M | 8 | 0.3 | 4.3 |  |  |
| *Tadarida brasiliensis* | F | 5 | 1.2 | 6.2 | 180 | 40 |
|  | M | 6 | 1.2 | 6.2 |  |  |

<sup>a</sup> Sex-biased sites were defined using the full model (beta ~ sex + age) with an FDR cutoff 0.05.
<sup>b</sup> Age-associated sites which showed evidence of age-associated variation (Punadj. < 0.05) in both sexes, with consistent sign, in sex-specific regression (beta ~ age).

**Figure S1.**
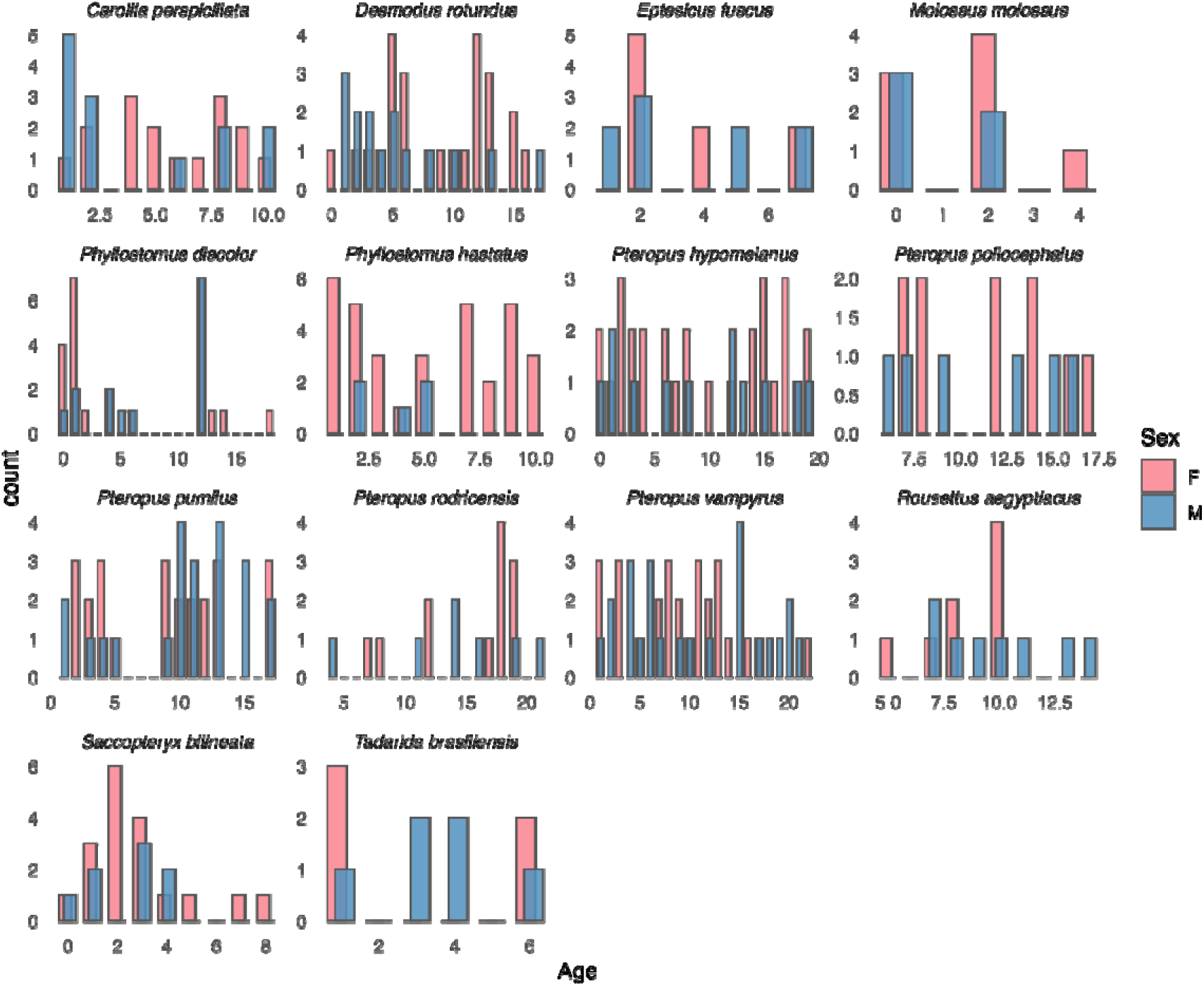
Numbers of samples of each sex, for each species, and the age distribution.

**Figure S2.**
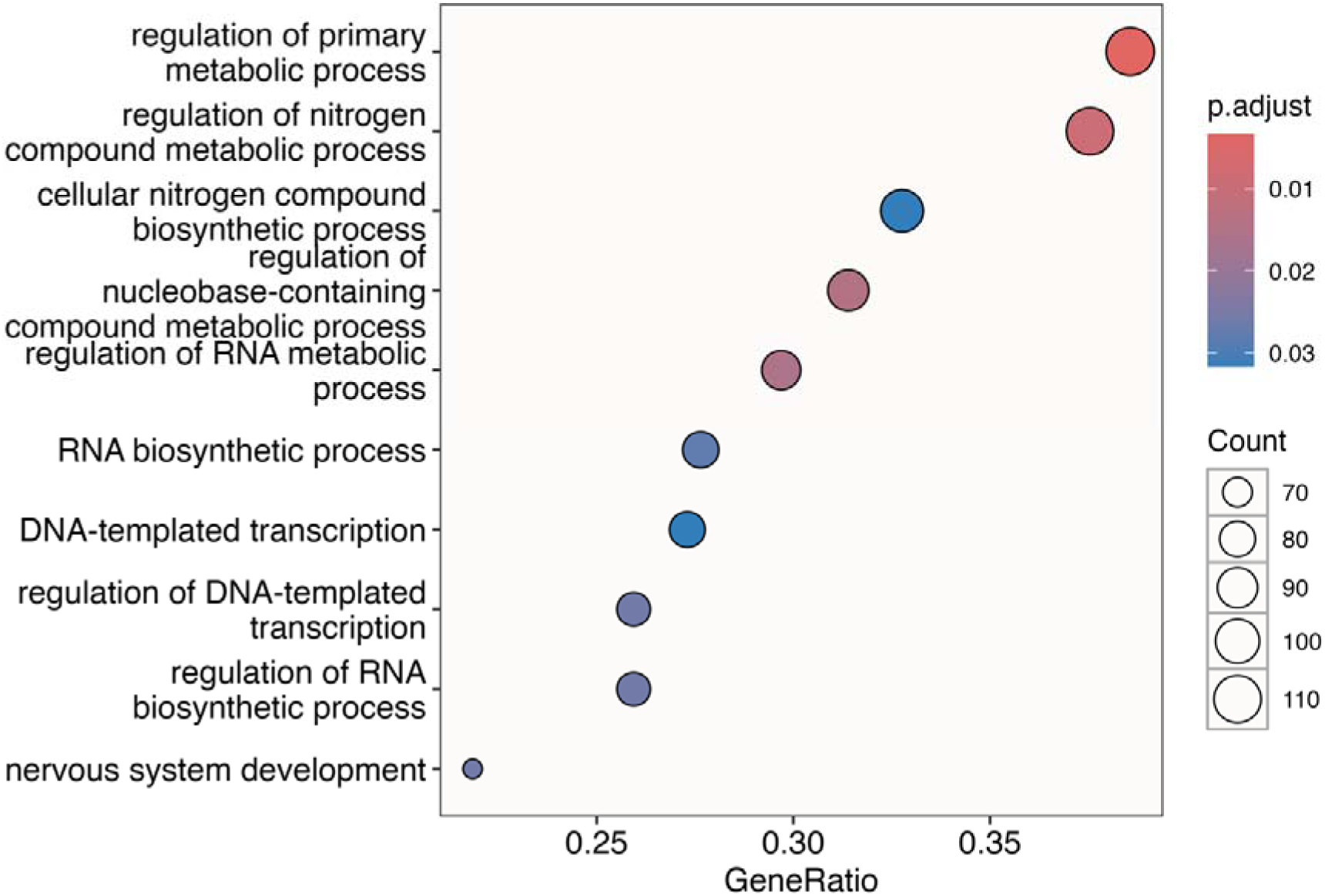
Overrepresented gene ontology terms among genes annotated for proximity to sites that showed sex-biased methylation in any species.

**Figure S3.**
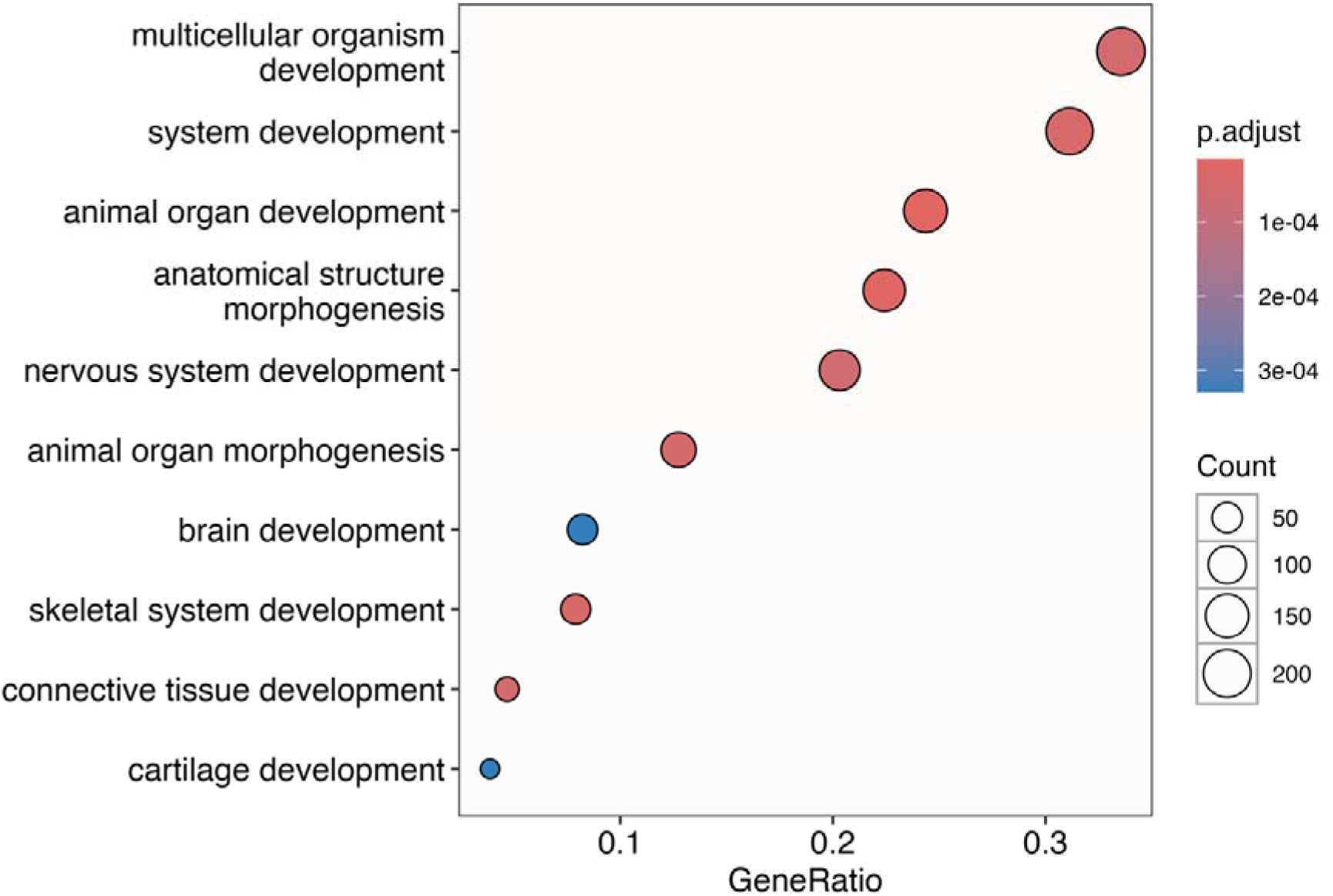
Overrepresented gene ontology terms among genes annotated for proximity to sites that showed age-associated methylation in any species.

**Fig S4.**
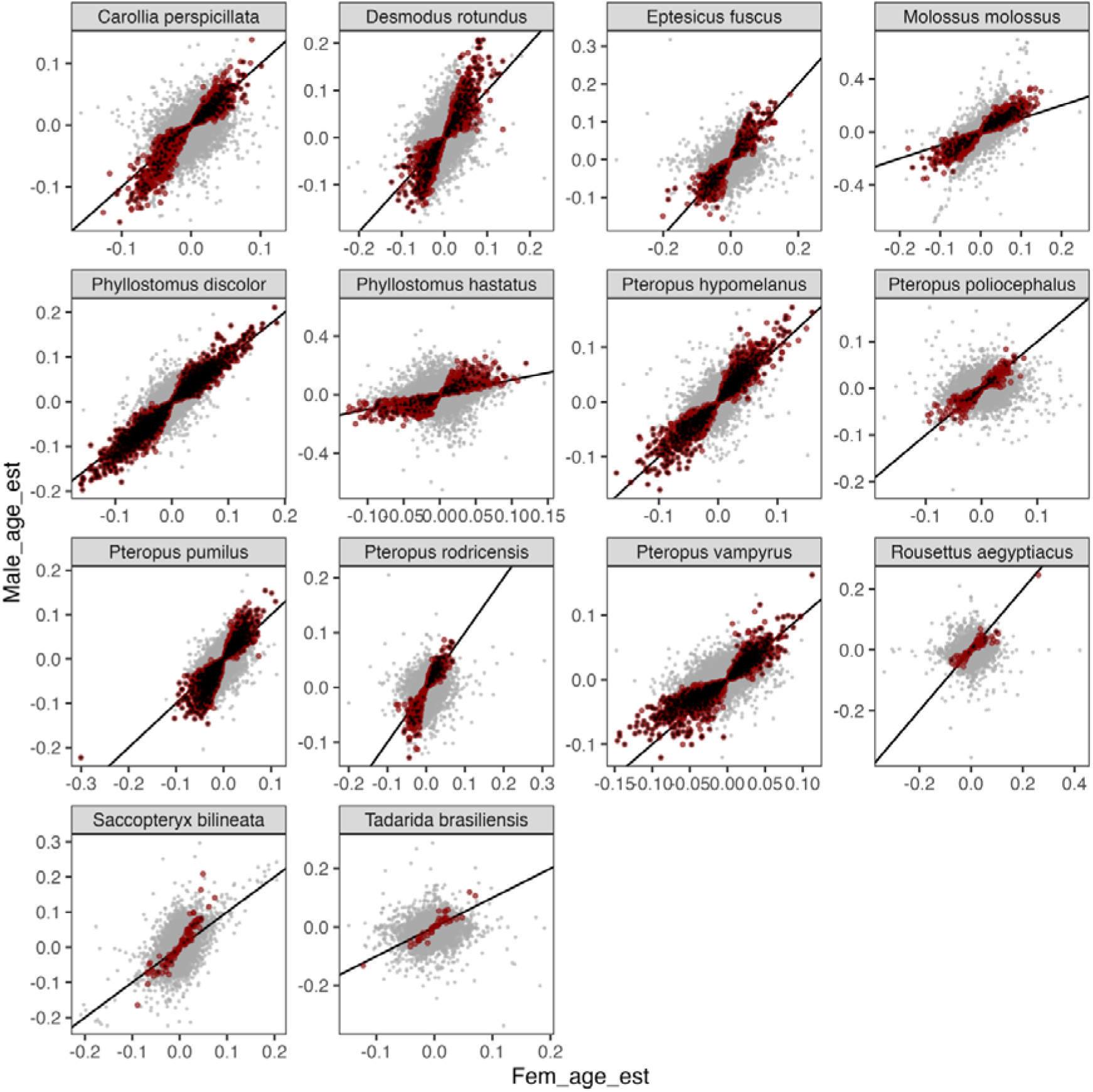
Correlations between male and female rates of age-associated slopes from sex-specific models of beta ∼ age. Red points show sites used in the analysis comparing slopes between the two. Black points show sites from the same analysis but more stringent P-value filtering (P_unadj._ < 0.01 in both sexes). Black lines show an intercept of 0 and slope of 1.

**Figure S5.**
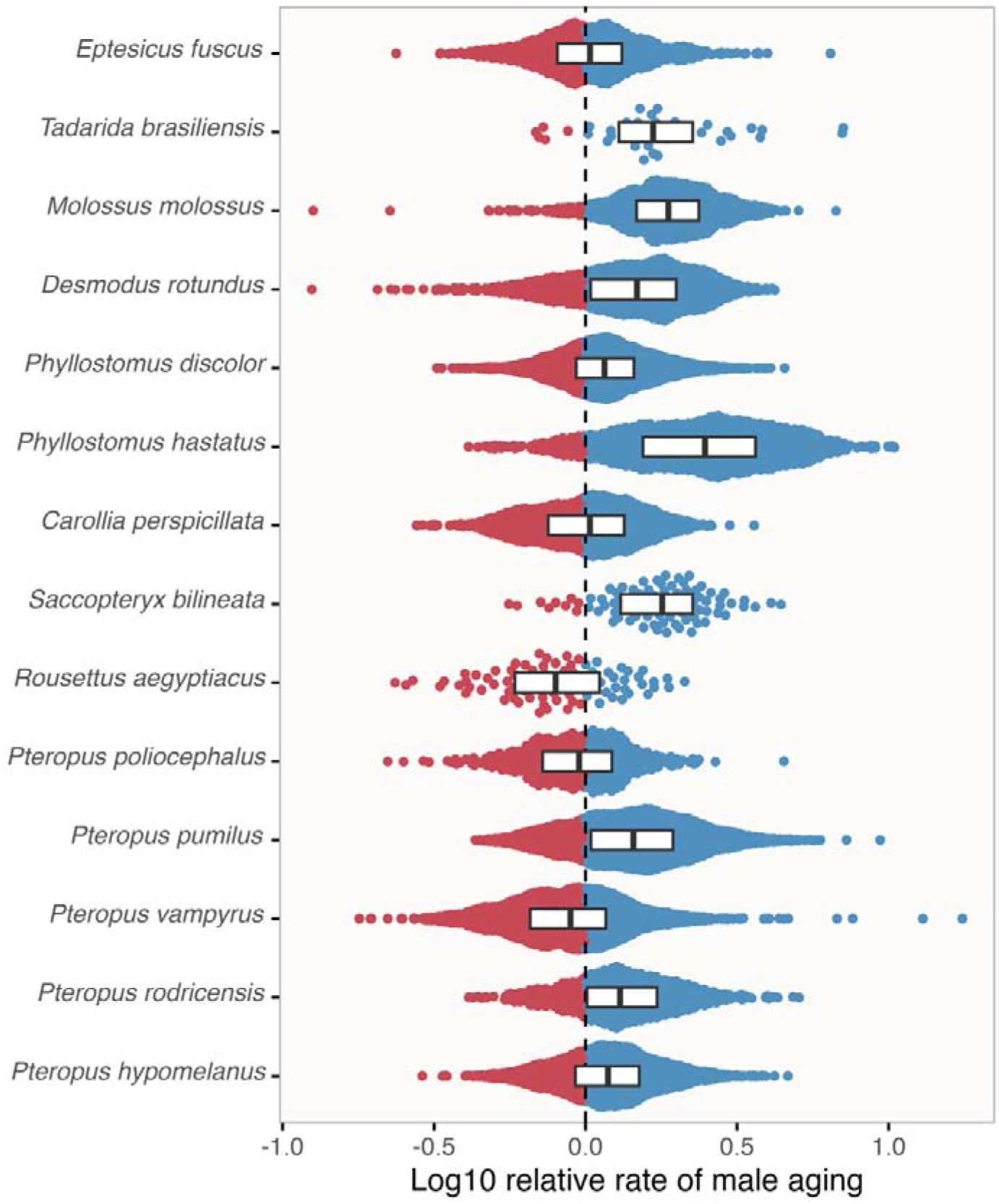
Sex-biased rates of change at age-associated CpG sites (autosomal sites). Each point illustrates relative rates of change in the proportion of methylated CpG sequences with age among sites that showed evidence of correlated age-associated change (P_unadj._ < 0.05) in both sexes in each species. Note the log10 scale, such that values greater-than or less-than 0 represent male and female-biased slopes of age-associated variation, respectively.

**Figure S6.**
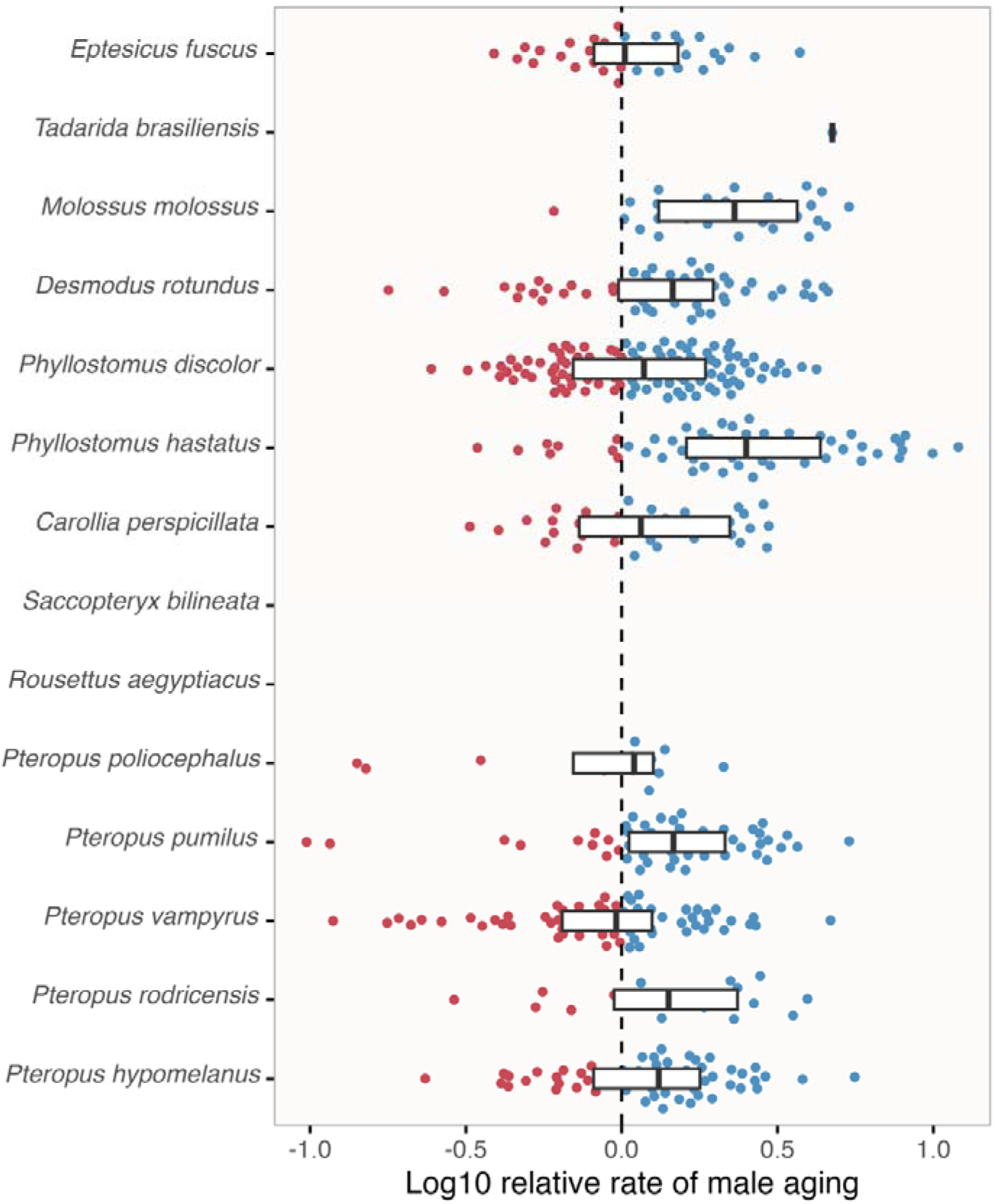
Sex-biased rates of change at age-associated CpG sites (X-linked sites). Each point illustrates relative rates of change in the proportion of methylated CpG sequences with age among sites that showed evidence of correlated age-associated change (P_unadj._ < 0.05) in both sexes in each species. Note the log10 scale, such that values greater-than or less-than 0 represent male and female-biased slopes of age-associated variation, respectively.

**Figure S7.**
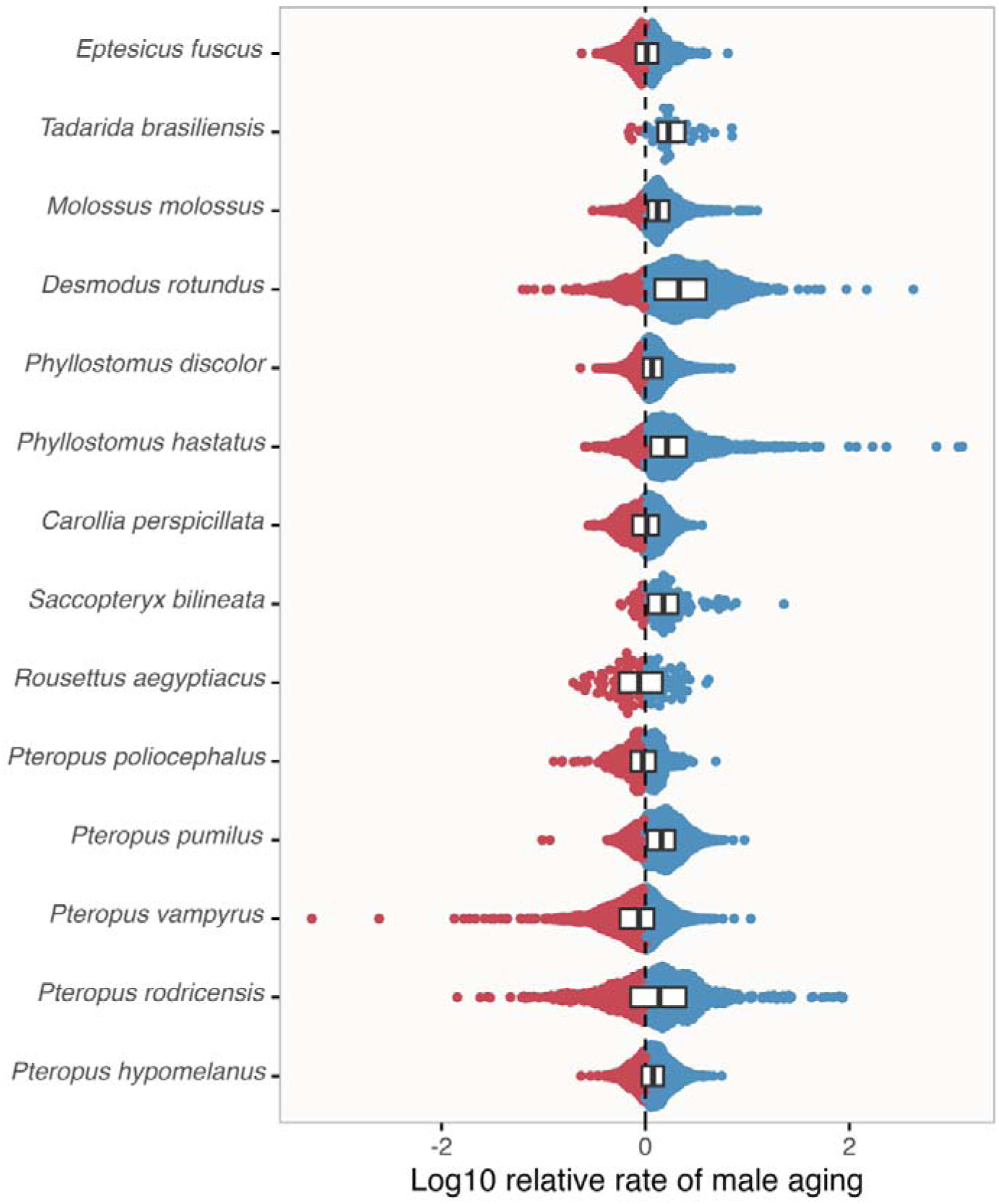
Sex-biased rates of change at age-associated CpG sites (truncated analysis). Each point illustrates relative rates of change in the proportion of methylated CpG sequences with age among sites that showed evidence of correlated age-associated change (P_unadj._ < 0.05) in both sexes in each species. Note the log10 scale, such that values greater-than or less-than 0 represent male and female-biased slopes of age-associated variation, respectively.

**Figure S8.**
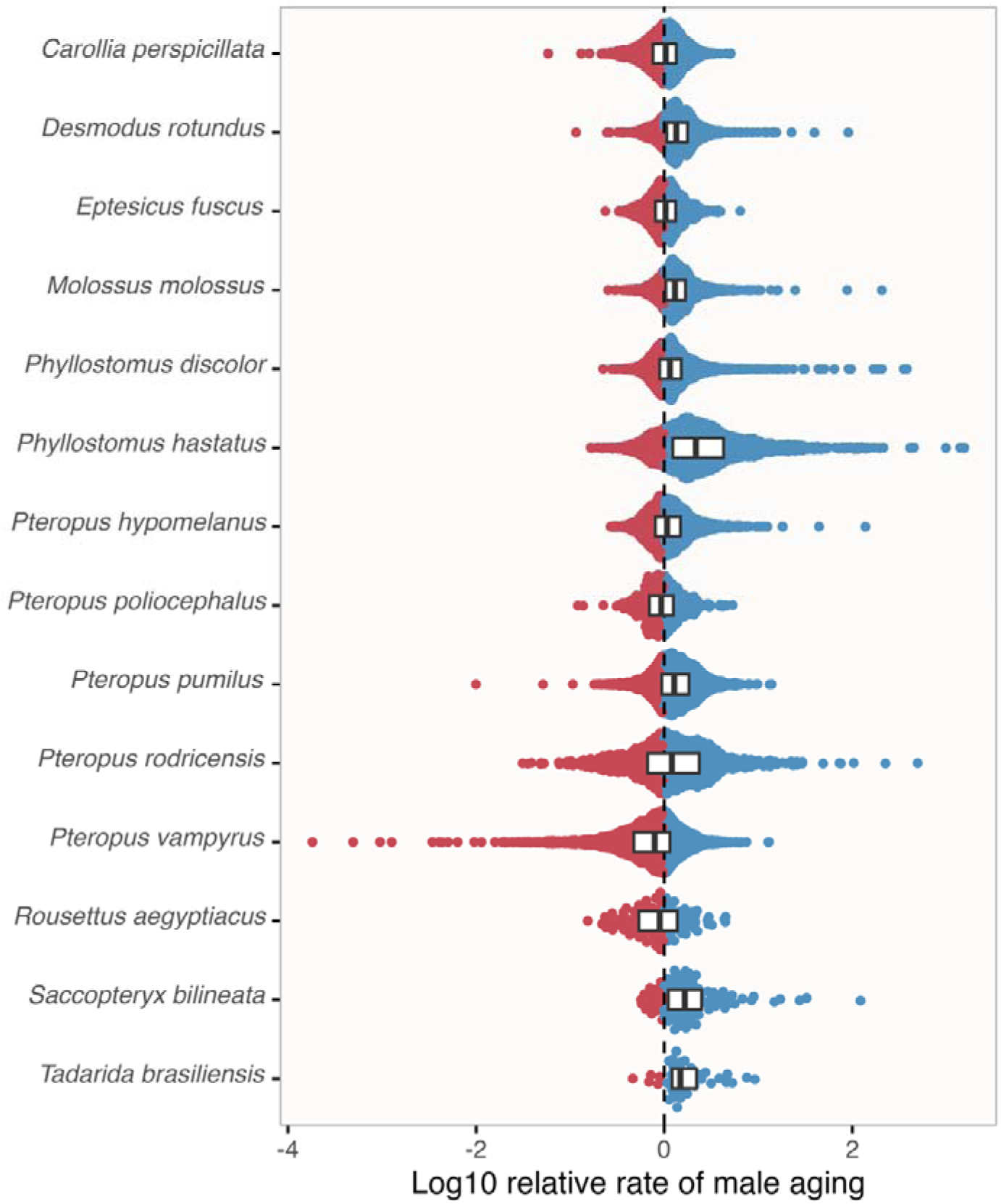
Sex-biased rates of change at age-associated CpG sites (paired analysis). Each point illustrates relative rates of change in the proportion of methylated CpG sequences with age among sites that showed evidence of correlated age-associated change (P_unadj._ < 0.05) in both sexes in each species. Note the log10 scale, such that values greater-than or less-than 0 represent male and female-biased slopes of age-associated variation, respectively.

**Figure S9.**
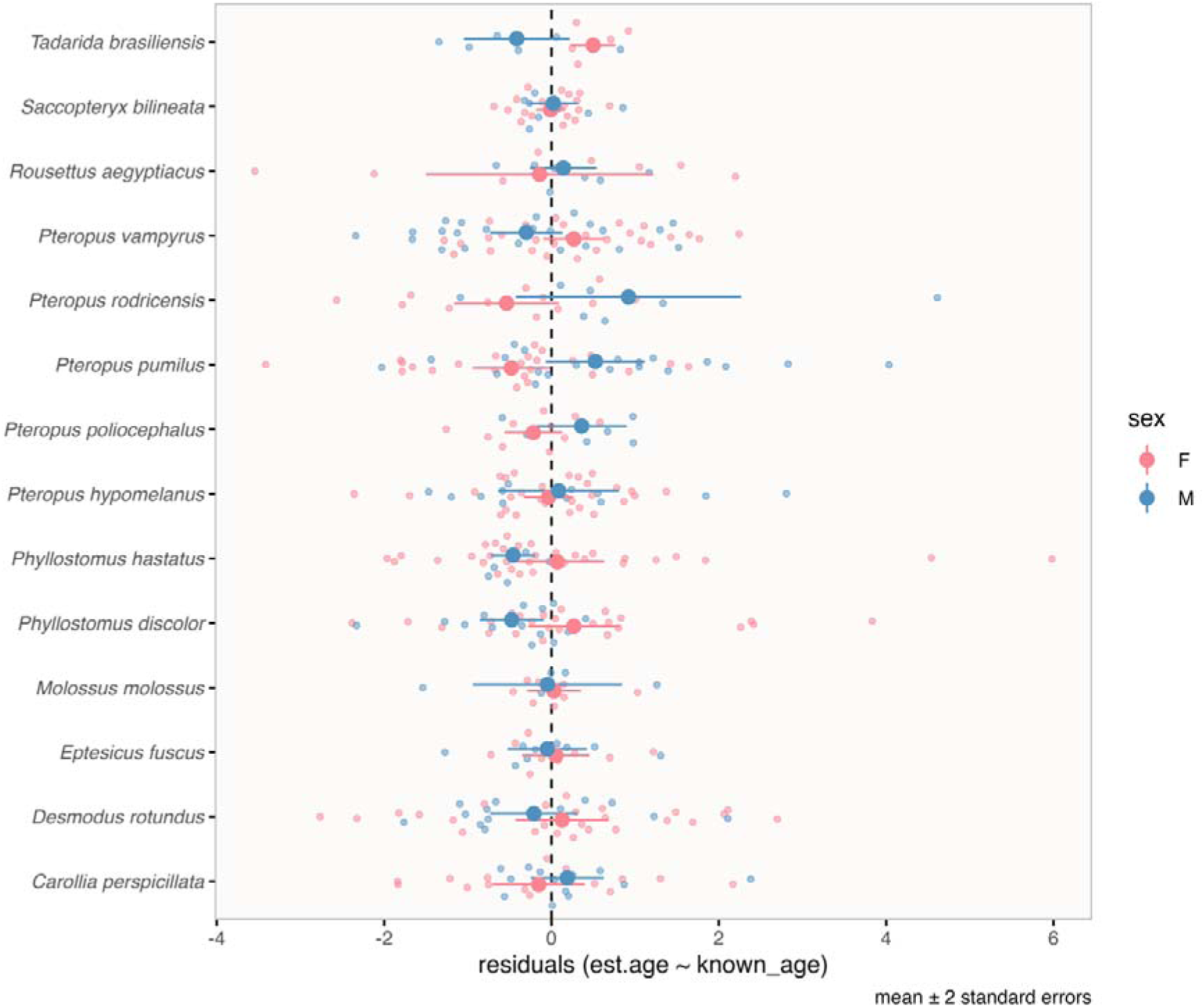
Residuals from regression of known chronological age on estimated age based on the ‘all bat’ clock^5^. Points show means and error bars shown 2 standard errors.

**Figure S10.**
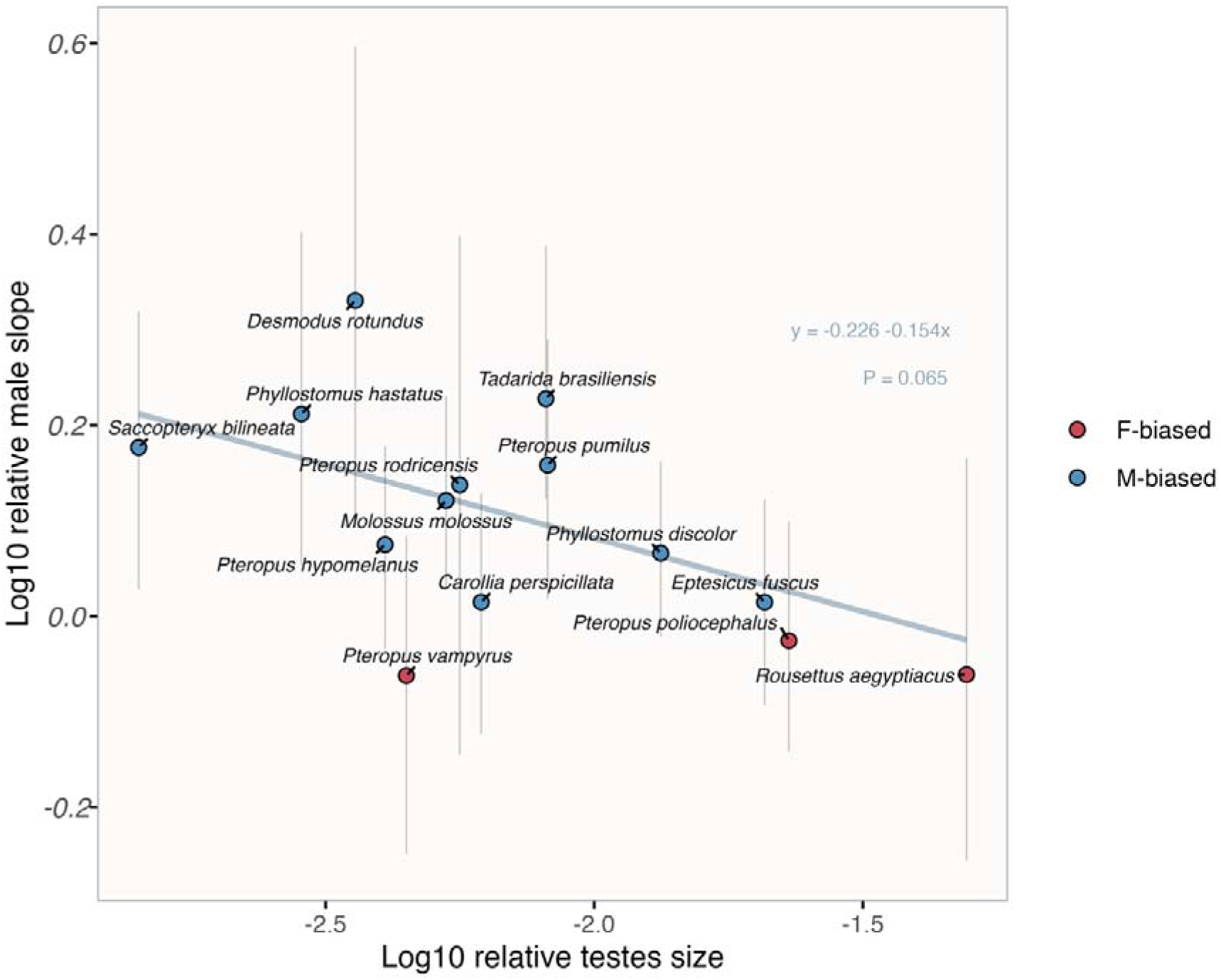
Relationship between relative testes mass and relative rates of methylation change in males (truncated analysis). Points and vertical lines illustrate median values and interquartile ranges with respect to relative male slopes of age-associated variation, also shown in Fig. 3. Blue and red points represent species in which males and females tended to have stronger slopes of age-associated variation, respectively. The solid line visualises the relationship predicted by phylogenetic generalized least squares regression.

**Figure S11.**
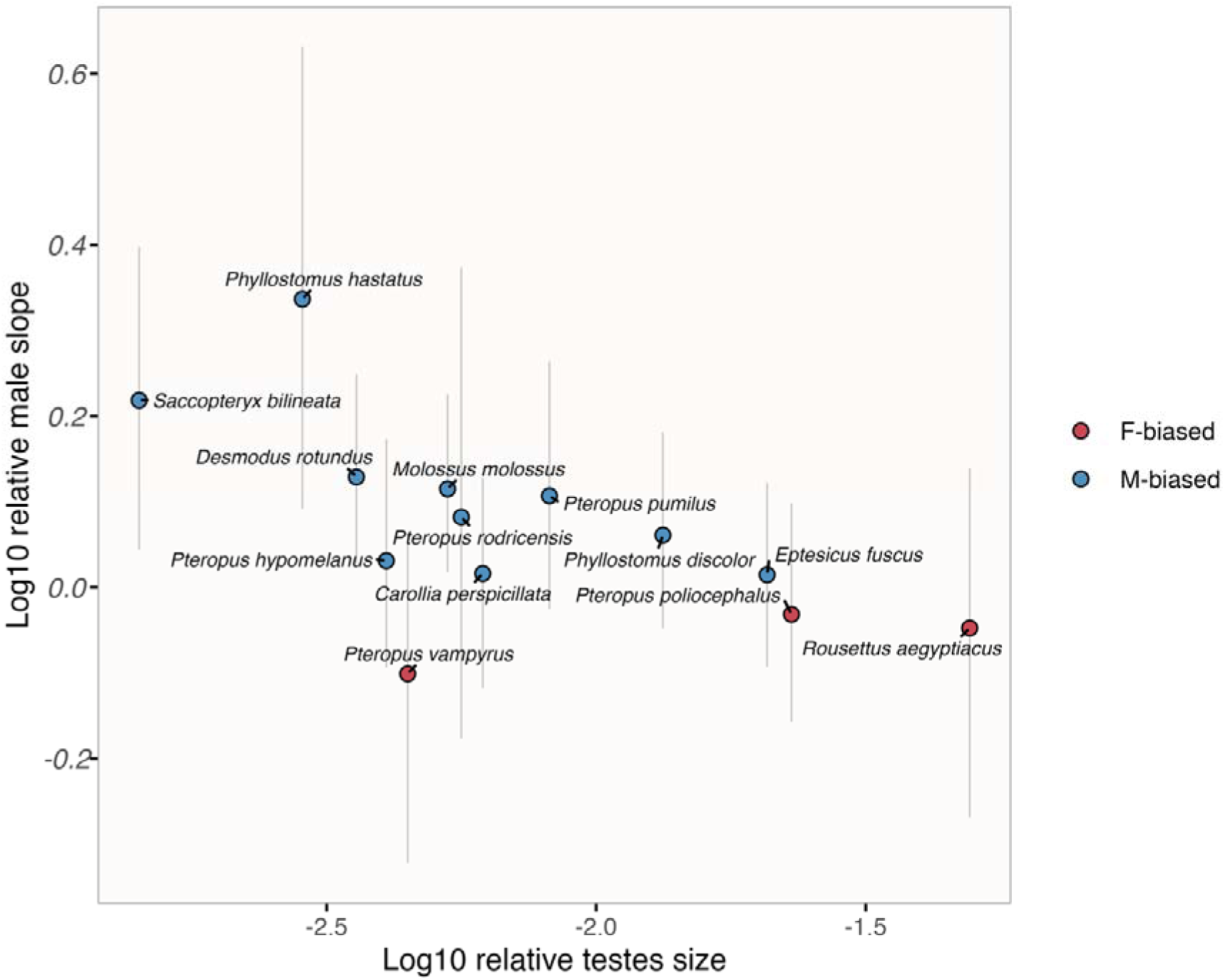
Relationship between relative testes mass and relative rates of methylation change in males (paired analysis). Points and vertical lines illustrate median values and interquartile ranges with respect to relative male slopes of age-associated variation, also shown in Fig. 3. Blue and red points represent species in which males and females tended to have stronger slopes of age-associated variation, respectively. Note that the regression line is not displayed as the model did not satisfy assumptions of PGLS due to non-normally distributed residuals.

